# Leveraging population-based clinical quantitative phenotyping for drug repositioning

**DOI:** 10.1101/130799

**Authors:** Adam S Brown, Danielle Rasooly, Chirag J Patel

**Affiliations:** Department of Biomedical Informatics, Harvard Medical School, Boston, MA 02115; Department of Biomedical Informatics, Harvard Medical School, 10 Shattuck St. Boston, MA 02115.

## Abstract

Computational drug repositioning methods can scalably nominate approved drugs for new diseases, with reduced risk of unforeseen side effects. The majority of methods eschew individual-level phenotypes despite the promise of biomarker-driven repositioning. In this study, we propose a framework for discovering serendipitous interactions between drugs and routine clinical phenotypes in cross-sectional observational studies. Key to our strategy is the use of a healthy and non-diabetic population derived from the National Health and Nutrition Examination Survey, mitigating risk for confounding by indication. We combine complementary diagnostic phenotypes (fasting glucose and glucose response) and associate them with prescription drug usage. We then sought confirmation of phenotype-drug associations in un-identifiable member claims data from Aetna using a retrospective self-controlled case analysis approach. We identify bupropion hydrochloride as a plausible antidiabetic agent, suggesting that surveying otherwise healthy individuals cross-sectional studies can discover new drug repositioning hypotheses that have applicability to longitudinal clinical practice.

## INTRODUCTION

Identifying new indications for previously approved drugs, known as drug repositioning, is an attractive alternative to the traditional drug discovery paradigm as previously approved drugs have substantially lower risk of unforeseen adverse events^1^. Computational drug repositioning builds on this premise by pre-screening for promising repositioning candidates, with current methods primarily relying on molecular data^2–5^ and/or the biological literature^6–10^. While these methods have been successful in predicting plausible repositioning candidates, a key challenge in computational repositioning is to provide direct evidence of candidate efficacy in humans, rather than relying on surrogate biomarkers or indirect evidence.

One alternative is to consider a single or a few quantitative phenotypes’ association with drug prescription history. In doing so, one can not only be certain that the phenotypes chosen are clinically relevant to a disease of interest, but also readily access effect sizes for power considerations in future clinical studies. While such a strategy is appealing, even a study limited to a single disease may be confounded due to shared disease etiology^11^, off-label drug usage^12^, and variable effects of drugs due to disease severity. We propose a novel framework in which the association between drugs and quantitative phenotypes is assessed in a non-institutionalized population who do not have the target disease for repositioning.

To demonstrate the potential of this strategy, we search for putative modulators of glycemic health in a normoglycemic and US-representative population of participants from the 2005-2012 National Health and Nutrition Examination Survey (NHANES). We evaluated associations between 134 prevalent drugs and two diabetes diagnostic phenotypes, fasting blood glucose and glucose after following a 2-hour oral glucose tolerance challenge (or, glucose response). By combining findings from two glycemic phenotypes, we identified a single potential antidiabetic candidate associated with lower glycemic phenotypes, the antidepressant bupropion. Notably, other commonly used antidepressants did not show multimodal antidiabetic potential. To replicate the association, we designed a retrospective self-controlled study in a normoglycemic cohort derived from un-identifiable member claims data provided by Aetna, and again verified that bupropion, but not other commonly prescribed antidepressants, was associated with lower levels of fasting blood glucose after exposure to the drug.

## RESULTS

### Association of bupropion with complementary diabetes phenotypes in a normoglycemic population

After excluding participants with a reported history of diabetes, abnormal fasting blood glucose (including diabetes and prediabetes, according to American Diabetes Association guidelines, >= 100 mg/dL), and who were currently prescribed an antidiabetic drug, we obtained a final NHANES-derived cross-sectional cohort size of 5,371 (see Table 1 for demographic characteristics of the cohort). Using the normoglycemic cohort, we performed comprehensive association testing between prescription drug use and either fasting blood glucose or blood glucose following an oral glucose tolerance test, adjusting for age, sex, race, and body mass index. Of the 134 prescription drugs with power (out of 1,133 total drugs tracked) to detect an association, the antidepressant bupropion was significantly and negatively associated with both glycemic phenotypes (survey-correct multivariate linear regression β < 0, FDR < 0.2, Figure 1A). Bupropion was associated with -2.5 mg/dL (95% CI: [-4.4, -0.6]) lower fasting blood glucose, and -10.4 mg/dL (95% CI: [-16.8, -4]) lower blood glucose following an oral glucose tolerance test (see Table S1 for demographic characteristics of the bupropion exposed subset). We also tested the association between both phenotypes and two commonly used antidepressants, duloxetine and escitalopram, with sufficient populations of exposed individuals. Neither duloxetine or escitalopram had significant effects on both glycemic phenotypes, suggesting that the association with bupropion is specific (see Table 2 for antidepressant results, and Table S2 for a full list of drugs with one significant glycemic association).

**Table 1.**
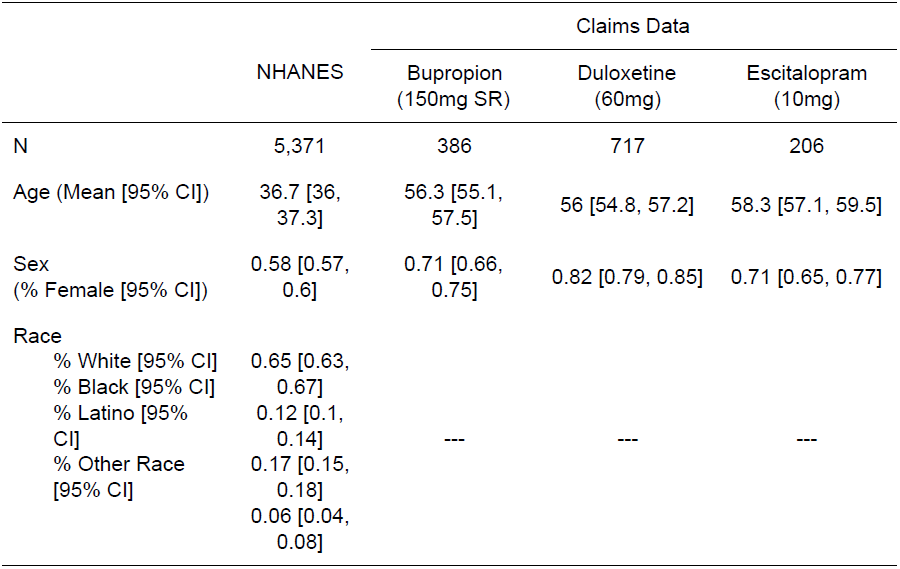
Demographic breakdown of NHANES (cross-sectional) and un-identifiable member claims data from Aetna (longitudinal) cohorts

|  | NHANES | Claims Data |  |  |
| --- | --- | --- | --- | --- |
|  |  | Bupropion<br>(150mg SR) | Duloxetine<br>(60mg) | Escitalopram<br>(10mg) |
| N | 5,371 | 386 | 717 | 206 |
| Age (Mean [95% CI]) | 36.7 [36, 37.3] | 56.3 [55.1, 57.5] | 56 [54.8, 57.2] | 58.3 [57.1, 59.5] |
| Sex<br>(% Female [95% CI]) | 0.58 [0.57, 0.6] | 0.71 [0.66, 0.75] | 0.82 [0.79, 0.85] | 0.71 [0.65, 0.77] |
| Race |  |  |  |  |
| % White [95% CI] | 0.65 [0.63, 0.67] |  |  |  |
| % Black [95% CI] | 0.12 [0.1, 0.14] | --- | --- | --- |
| % Latino [95% CI] | 0.17 [0.15, 0.18] |  |  |  |
| % Other Race [95% CI] | 0.06 [0.04, 0.08] |  |  |  |

**Table 2.** Antidepressant association with diabetes health in NHANES (cross-sectional)

| Drug Name | Participants Prescribed<br>(of 5,371) | Observed effect on<br>fasting glucose<br>(mg/dL)<br>[95% CI] | FDR | Observed effect on<br>glucose tolerance<br>(mg/dL)<br>[95% CI] | FDR |
| --- | --- | --- | --- | --- | --- |
| **Bupropion** | **38** | **-2.5 [-4.4, -0.6]** | **0.12** | **-10.4 [-16.8, -4]** | **0.07** |
| Escitalopram | 43 | -2 [-3.8, -0.1] | 0.26 | -13.6 [-22.4, -4.9] | 0.08 |
| Duloxetine | 24 | -1 [-4, 3] | 0.84 | -8 [-21, 6] | 0.63 |

**Figure 1.**
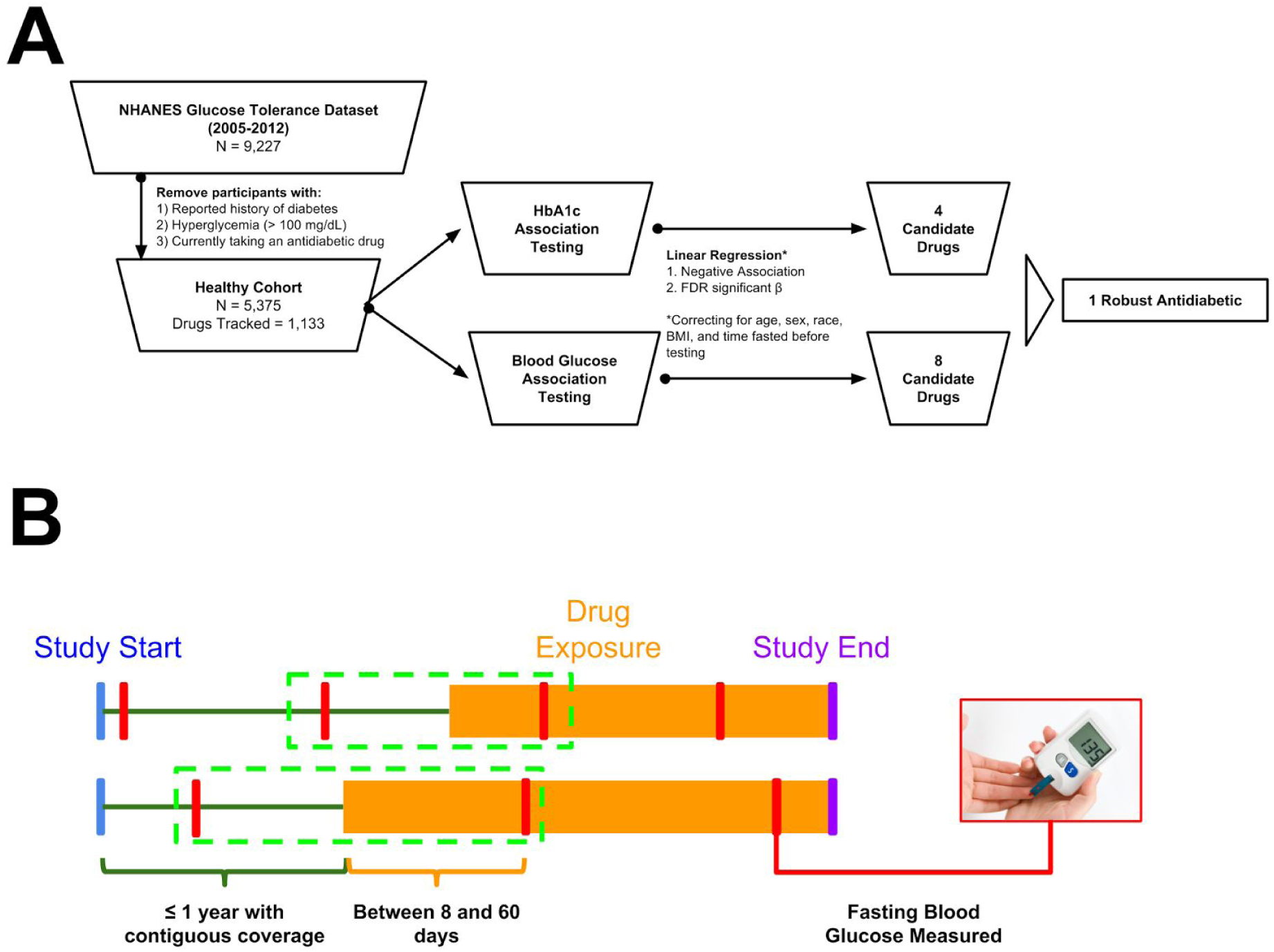
Quantitative-phenotype based repositioning overview. A) NHANES (cross-sectional) quantitative-phenotype based repositioning workflow. B) Conceptual diagram of claims data-based replication efforts.

### Replication in a retrospective self-controlled study

To replicate the association of bupropion with improved fasting blood glucose, we performed a self-controlled study of fasting blood glucose using un-identifiable member claims data from the Aetna Insurance Company, containing information from 50 million individuals over 9 years. We extracted three non-diabetic populations with exposure either to bupropion or to one of two control antidepressants, duloxetine, and escitalopram (see Table 1 for demographic characteristics of each drug-exposed cohort). Age, sex, and race were not significantly different between drug-exposed cohorts (ANOVA, p > 0.1 for age, sex, and race). Within each drug-exposed cohort, we identified individuals with a fasting glucose measurement up to a year before being exposed (glucose measurements are typically performed on an annual basis^13^), and within two months after being exposed (with a buffer of 8 days to reach steady-state drug concentration). Within each drug, we selected dosage forms with at least 198 individuals for well-powered association testing. For bupropion, the only dosage form with sufficient individuals was 150mg sustained release with 383 individuals (of 11 total dosage forms), for duloxetine only 60mg was powered with 717 individuals (of 5 total dosage forms), and for escitalopram only 10mg was powered with 206 individuals (of 3 total dosage forms). We found that only bupropion 150mg sustained release was associated with significantly decreased fasting blood glucose (mean difference -1.88 mg/dL, 95% CI: [-2.91, -0.85], p < 0.001, see Table 3).

**Table 3.**
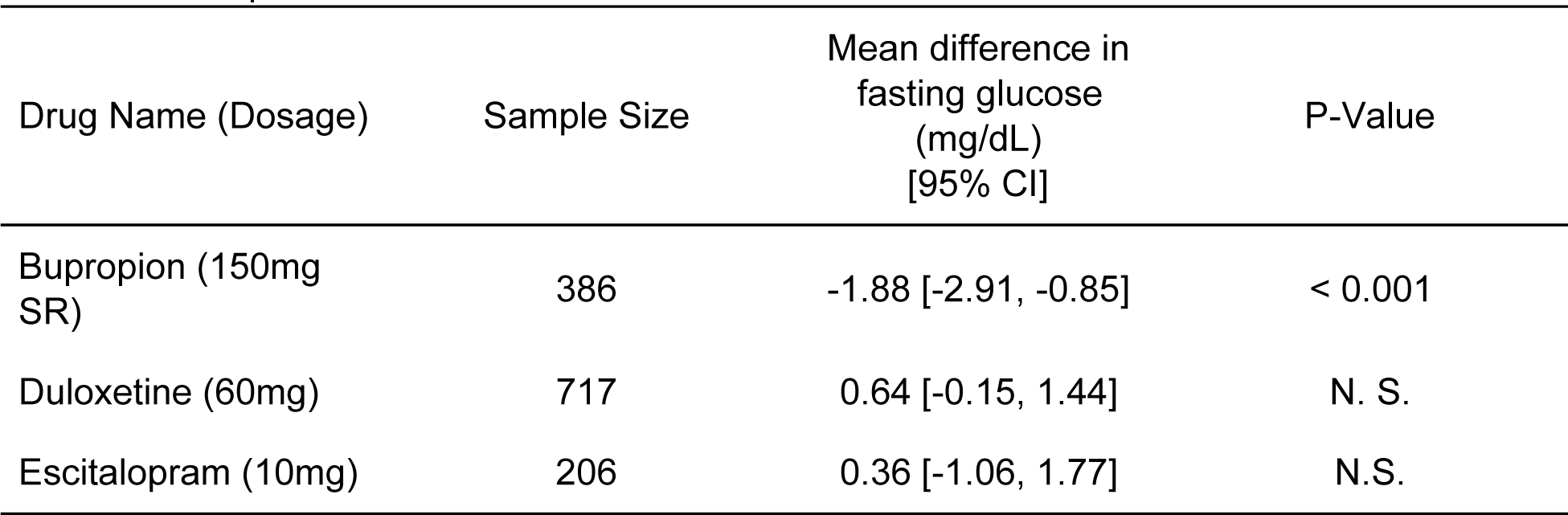
Un-identifiable member claims data from Aetna (longitudinal) confirmation analysis of selected antidepressants

## DISCUSSION

In this study, we describe preliminary results from a novel quantitative phenotype-based drug repositioning methodology. Our methodology uses a combination of complementary quantitative phenotypes to efficiently reduce the number of potential repositioning candidates. Our method enables straightforward follow up in prospective investigations, and provides estimation of the population sizes required to detect modulation of disease-relevant phenotypes by a candidate drug. Furthermore, we perform all association testing in a non-diseased and otherwise nation-wide representative population to avoid common sources of confounding.

To demonstrate our complementary phenotype-based approach, we predicted repositioning candidates with the potential to modulate diabetes health using a 5,137 person non-diabetic cohort in NHANES. Fasting blood glucose and glucose following an oral glucose tolerance test capture related, but distinct etiological components of diabetes health, and impairment in either test implies distinct disease etiology (hepatic and muscular insulin resistance respectively).^14^ By combining these two phenotypes we identified a single candidate, the antidepressant bupropion, associated with improved glycemic status across multiple etiological pathways in diabetes.

Bupropion is well-known as a treatment for obesity comorbid with diabetes, both alone^15,16^ and in combination with naltrexone^17^, as well as a monotherapy for comorbid depression and diabetes^18^. What is unclear from the previous studies, however, is the degree to which the observed effects were caused by improvement in body mass index (BMI) or depression, which subsequently led to improvement in glycemic status (confounding by indication). In contrast, we explicitly address confounding by adiposity, depression status, and glycemic status: (1) by explicitly adjusting for BMI in all associations, (2) by testing other commonly used antidepressants associations with improved glycemic health, as measured by fasting glucose and glucose response, and (3) by testing for associations in a non-diabetic and normoglycemic cohort, decreasing the likelihood of more complex confounding scenarios, such as a single upstream process controlling both depression status and diabetic health (e.g. statins lower low-density lipoprotein levels, which in turn lowers heart attack and stroke risk simultaneously). In the future, we hope to follow up on the promise of bupropion as a multimodal antidiabetic, and propose clinical studies to understand its efficacy not only in healthy participants, but also in pre-diabetics and diabetics, alone and in combination with the current standard of care.

Another key benefit of our method is the ability to design clinical studies derived from our potential discoveries; we demonstrated this benefit by performing a retrospective self-controlled study using un-identifiable member claims data from Aetna. By designing an experiment by which patients serve as their own control, we avoid time-invariant confounding^19^. We successfully replicated that bupropion alone among the antidepressants considered is associated with lower fasting blood glucose. Despite escitalopram’s significant improvement of glucose response in the NHANES cohort, we did not observe a significant impact on fasting blood glucose in the self-controlled study. This result underscores the importance of combining multiple quantitative phenotypes to achieve high specificity in repositioning candidates. Furthermore, to enable future clinical trials, quantitative phenotype-driven studies should carefully select phenotypes with the goal of not only identifying repositioning candidates with high specificity, but also with clinical study design in mind, considering both clinical relevance and statistical power.

While we have discussed the potential of combining complementary quantitative phenotypes for drug repositioning, we do note that it has some limitations. Chief among these is the requirement that non-diseased individuals are assayed for quantitative phenotypes. While common diseases by necessity have associated routine diagnostics, for example fasting blood glucose and glucose tolerance for diabetes or lipid levels and blood pressure for cardiovascular disease, repositioning for rarer diseases may require non-standard tests. We expect that this limitation will diminish over time with the development of birth cohorts (e.g. ALSPAC ^20^ among others), and large biobanking initiative (e.g. UK Biobank ^21^ among others), most of which include clinical phenotyping of all participants to complement a variety of ‘omic measurements. A second potential limitation is the requirement for multiple complementary quantitative phenotypes that are associated with the disease of interest for repositioning. For diseases where such phenotypes are not available, further biomarker identification may be required before using our repositioning strategy. We note that any quantitative phenotype-based methodology would likely require disease-associated phenotypes before producing meaningful and interpretable results. Lastly, because we assess all associations between drugs and phenotypes in a non-diseased population, it will be important to verify any repositioning candidates that arise from this method in disease sufferers.

## STUDY HIGHLIGHTS

**What is the current knowledge on the topic?** Observationalstudies are emerging as ways to search for repositioning candidates, yet are fraught with bias and do not consider quantitative phenotypes.

**What question did this study address?** Is it possible to use health monitoring surveys and longitudinal administrative population databases coupled with continuous phenotypes to search for and replicate new repositioning candidates?

**What this study adds to our knowledge?** We present a novel approach for drug repositioning that harnesses health monitoring surveys and multiple clinical trait phenotypes to avoid confounding bias and increase specificity of evidence for repositioning discovery.

**How might this change drug discovery, development, and/or therapeutics?** Our method enhances the repositioning process using quantitative phenotyping from humans, poteally closing the gap between computational and clinical drug repositioning.

## METHODS

### Cross-sectional study cohort

The cross-sectional study cohort was derived from a combination of four independent waves of the continuous National Health and Nutrition Examination Survey (NHANES): the 2005-2006, 2007-2008, 2009-2010, and the 2011-2012 surveys^22^. NHANES is a cross-sectional survey conducted by the United States Centers for Disease Control and Prevention (US CDC), wherein a large number of participants are recruited to answer a number of questions pertaining to their medical, psychosocial, and sociodemographic histories. A subset of respondents also receive extensive anthropometric and laboratory testing, including a variety of routine clinical measures. For this study, several variables were obtained for each respondent, including: (1) self-reported history of diabetes (field DIQ010 from the respective DIQ questionnaire datasets), (2) fasting blood glucose and fasting time, as well as glucose taken at 2-hours post oral glucose tolerance test (field LBXGLU and PHASFSTHR, LBXGLT respectively from the respective GLU laboratory datasets) (3) self-reported prescription drug usage at the time of interview (including generic drug names and drug category, as defined in the Lexicon Plus^®^ database, Cerner Multum, Inc, see Figure 1A). Respondents without completed information for any of these fields were excluded from further analysis. To obtain a non-diabetic final cohort for association testing, respondents were filtered to include those with no reported history of diabetes, no use of antidiabetic medications at the time of interview, and normal glycemic status (fasting blood glucose less than 100 mg/dL according to American Diabetes Association guidelines^13^).

### Drug-phenotype association in the cross-sectional study cohort

Multivariate linear regression was performed to individually test the association of usage of 1,133 drugs (1.16 [1.11, 1.22], mean [95%CI] drugs per participant) and either fasting blood glucose or blood glucose taken at 2-hours post oral glucose tolerance test (controlling for age, sex, race, body mass index [BMI]), and number of hours fasted). Regression coefficients and significance were estimated using the *survey* package in *R* to account for the stratified design of NHANES^23^. To avoid erroneous associations, drugs with less than 12 prescribed individuals were removed from further analysis (see^24^ for rationale). For the remaining 134 drugs, regression coefficients and significance were obtained and corrected for multiple testing using the False Discovery Rate (FDR) method^25^. Drugs with significant (FDR < 0.2), negative associations with both fasting blood glucose and blood glucose following a 2-hour post oral glucose tolerance test were considered candidate antidiabetic agents (Figure 1A).

### Longitudinal validation cohort and self-controlled analyses

The longitudinal validation cohorts were derived from un-identifiable member claims data from Aetna, spanning 8 years (2008-2016) with over 50 million lives in all 50 states. To obtain drug-specific non-diabetic cohorts the following restrictions were made: (1) patients did not have any instance of diabetes diagnosis (ICD-9 codes 250-250.93), (2) patients were prescribed one of the study drugs (bupropion, duloxetine, or escitalopram), (3) patients were enrolled in an insurance plan for at least one year before the first prescription date for the respective study drug, (4) patients had a fasting blood glucose measurement up to one year before starting the respective study drug (“pre” measurement), and between 8 and 60 days after starting (“post” measurement) (Figure 1B). Timepoints were chosen to enable the collection of at least two glucose measurements (typically taken annually per American Diabetes Association recommendations), and for individuals to have reached steady-state concentrations of study drug (typically ∼1 week) while still minimizing time-dependent variation. Following cohort creation, self-controlled analysis was performed for each dose of each study drug using a paired t-test between the pre- and post-glucose measurements as previously described^19^. More recent methods have proposed additional corrections to the simple paired t-test method; however, these methods examine longer time periods of exposure (which allows for additional confounders to accumulate), and a population containing cases in addition to healthy controls^26^. In contrast, the method described here relies on a very short analysis window. Dosage forms of study drugs with fewer than 198 participants were removed from further analysis due to power considerations (assuming a small effect size, Cohen’s d equal to 0.2, requiring power of 80% or greater, see^27^ for details). Study drugs with significant (t-test p-value < 0.05) and negative associations (improved glucose response) with the pre- to post-drug regimes were considered replicated agents.

## ACKNOWLEDGEMENTS

ASB and DR were supported by an National Institutes of Health (NIH) Training grant from the National Human Genome Research Institute (NHGRI), T32HG002295-13. CJP is supported by a National Institute of Environmental Health Sciences (NIEHS) R00 ES023504, R21 ES025052, National Science Foundation Big Data Spoke (1636870), and a gift from Agilent Technologies.

## AUTHOR CONTRIBUTIONS

ASB and CJP conceived of the study. ASB and DR conducted all statistical analyses. ASB, DR, and CJP wrote the manuscript.

## TABLES

**Table S1.**
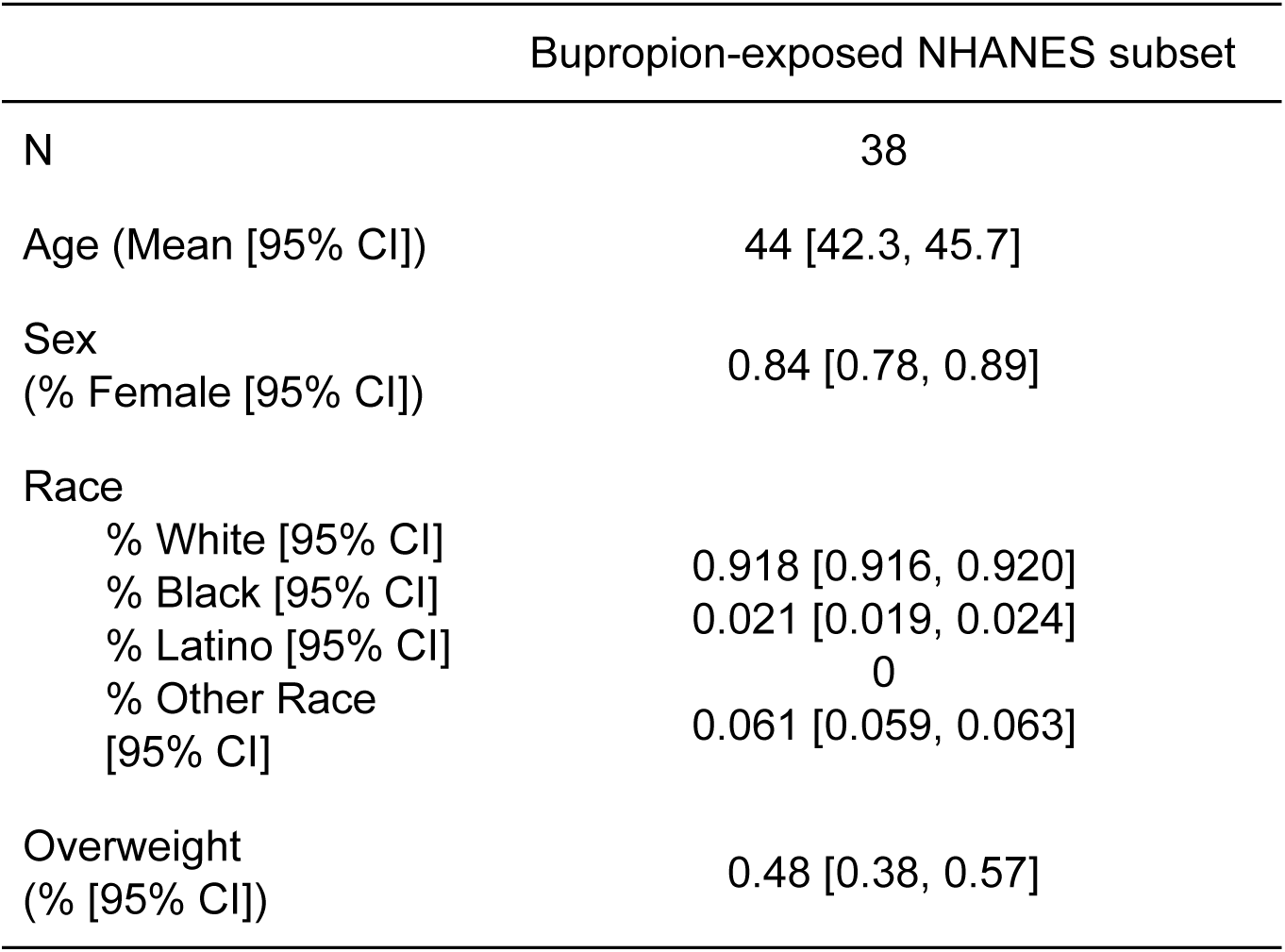
Demographic breakdown of bupropion-exposed NHANES (cross-sectional) participants

**Table S2.** NHANES (cross-sectional) results for drugs associated with decreased fasting glucose and glucose response

| Drug Name | Observed effect on<br>fasting glucose (mg/dL)<br>[95% CI] | FDR | Observed effect on<br>glucose tolerance (mg/dL)<br>[95% CI] | FDR |
| --- | --- | --- | --- | --- |
| **Bupropion** | **-2.5 [-4.4, -0.6]** | **0.12** | **-10.4 [-16.8, -4]** | **0.07** |
| Escitalopram | -2 [-3.8, -0.1] | 0.26 | -13.6 [-22.4, -4.9] | 0.08 |
| Phenylephrine | 0.3 [-1.6, 2.2] | 0.89 | -14.4 [-24.1, -4.8] | 0.09 |
| Setraline | 0.1 [-1.5, 1.8] | 0.91 | -9.7 [-15.1, -4.3] | 0.04 |
| Ethinyl<br>Estradiol | -1.8 [-2.9, -0.7] | 0.04 | 9.3 [4.1, 14.6] | 0.04 |
| Acyclovir | -7.2 [-10.9, -3.5] | 0.02 | -17.7 [-32.5, -2.8] | 0.21 |
| Levothyroxine | -1.2 [-2.2, -0.2] | 0.20 | -0.4 [-5.1, 4.4] | 0.96 |
| Levonorgestrel | -2 [-3.6, -0.4] | 0.15 | 8.3 [-2.2, 18.8] | 0.51 |
| Gabapentin | -2.5 [-4.4, -0.6] | 0.12 | -3.6 [-26.7, 19.5] | 0.94 |
| Norgestimate | -3.3 [-5.7, -0.9] | 0.12 | 6.7 [-5.5, 18.9] | 0.64 |
| Etonogestrel | -6.2 [-10.1, -2.3] | 0.06 | -9.4 [-18.3, -0.5] | 0.26 |

